# Heterogeneous brain region-specific responses to astrocytic mitochondrial DNA damage in mice

**DOI:** 10.1101/2024.05.29.596517

**Authors:** Daniela A. Ayala, Anthony Matarazzo, Bonnie L. Seaberg, Misha Patel, Eliana Tijerina, Camryn Matthews, Gabriel Bizi, Ashton Brown, Alan Ta, Mendell Rimer, Rahul Srinivasan

## Abstract

Astrocytes use Ca^2+^ signals to regulate multiple aspects of normal and pathological brain function. Astrocytes display context-specific diversity in their functions, and in their response to noxious stimuli between brain regions. Indeed, astrocytic mitochondria have emerged as key players in governing astrocytic functional heterogeneity, given their ability to dynamically adapt their morphology to regional demands on their ATP generation and Ca^2+^ buffering functions. Although there is reciprocal regulation between mitochondrial dynamics and mitochondrial Ca^2+^ signaling in astrocytes, the extent of this regulation into the rich diversity of astrocytes in different brain regions remains largely unexplored. Brain-wide, experimentally induced mitochondrial DNA (mtDNA) loss in astrocytes showed that mtDNA integrity is critical for proper astrocyte function, however, few insights into possible diverse responses to this noxious stimulus from astrocytes in different brain areas were reported in these experiments. To selectively damage mtDNA in astrocytes in a brain-region-specific manner, we developed a novel adeno-associated virus (AAV)-based tool, Mito-PstI, which expresses the restriction enzyme PstI, specifically in astrocytic mitochondria. Here, we applied Mito-PstI to two distinct brain regions, the dorsolateral striatum, and the hippocampal dentate gyrus, and we show that Mito-PstI can induce astrocytic mtDNA loss *in vivo*, but with remarkable brain-region-dependent differences on mitochondrial dynamics, spontaneous Ca^2+^ fluxes and astrocytic as well as microglial reactivity. Thus, AAV-Mito-PstI is a novel tool to explore the relationship between astrocytic mitochondrial network dynamics and astrocytic mitochondrial Ca^2+^ signaling in a brain-region-selective manner.

## INTRODUCTION

Astrocytes have emerged as critical participants in governing neuronal function during health and disease (Stogsdill, Harwell, & Goldman, 2023). Once thought of as a group of rather homogenous support cells, compelling evidence has revealed that protoplasmic astrocytes are heterogenous (Batiuk et al., 2020; John Lin et al., 2017) and display context-dependent heterogeneity in their functions and in their response to injurious stimuli within and across brain regions (Khakh & Deneen, 2019; Zimmer, Orr, & Orr, 2024).

Astrocytic heterogeneity between brain regions extends to important organelles within these cells, including mitochondria. Indeed, we have recently shown that astrocytic mitochondria in the hippocampus (HPC) and dorsolateral striatum (DLS) are morphologically and functionally distinct (Huntington and Srinivasan 2021). Astrocytic mitochondria are a particularly relevant organelle in terms of astrocytic heterogeneity because they have robust spontaneous Ca^2+^ fluxes that respond to neurotransmitter agonists (Bazargani & Attwell, 2016; Huntington & Srinivasan, 2021; Jackson & Robinson, 2018), a distinct functional proteomic profile (Fecher et al., 2019), and spatial segregation into somata versus processes with regard to Ca^2+^ fluxes and morphology (Huntington & Srinivasan, 2021; Jackson & Robinson, 2018). In addition, mitochondria are very dynamic organelles, constantly undergoing fission, fusion, and turnover. They specialize in the production of ATP by oxidative phosphorylation (OXPHOS), but are also capable of buffering intracellular Ca^2+^. Interestingly, there is reciprocal regulation between mitochondrial dynamics and mitochondrial Ca^2+^ signaling in astrocytes (Jackson & Robinson, 2018; Stephen, Gupta-Agarwal, & Kittler, 2014), however, to our knowledge, the extent of this regulation into the rich diversity of astrocytes in different brain regions remains largely unexplored.

In this study, we exploit mitochondrial targeting of bacterial restriction enzymes as an approach to manipulate the mitochondrial genome, giving rise to mitochondrial DNA (mtDNA) loss in a cell-type-specific manner (Bacman, Williams, Pinto, & Moraes, 2014). Thus, mitochondrially targeted PstI, which cuts at two sites on mouse mtDNA, has been expressed in skeletal muscle fibers and neurons *in vivo*, primarily resulting in mtDNA depletion, deletion by recombination, and chronic reduction of OXPHOS (Fukui & Moraes, 2009; Pickrell, Pinto, Hida, & Moraes, 2011; Srivastava & Moraes, 2005). Brain-wide astrocytic mtDNA depletion was previously achieved by conditional abrogation of *Twinkle*, a nuclear encoded helicase required for mtDNA replication (Ignatenko et al., 2018). Although this work demonstrated the absolute necessity of mtDNA integrity for proper astrocyte function, few insights into possible diverse responses to this noxious stimulus from astrocytes in different brain areas were reported in these experiments. To selectively damage mtDNA in astrocytes in a brain-region-specific fashion, we have developed a novel adeno-associated virus (AAV)-based tool, Mito-PstI, which expresses PstI, specifically in astrocytic mitochondria. Here, we have applied this tool to the dentate gyrus the HPC and the DLS, two well-studied areas of the brain, relevant for Parkinson’s and Alzheimer’s disease, respectively. Our choice of HPC and DLS as brain regions in this study stems from our previous report in which we demonstrate differences in spontaneous mitochondrial Ca^2+^ influx events in astrocytes from these two brain regions (Huntington & Srinivasan, 2021).

Our results show that Mito-PstI can induce astrocytic mtDNA loss *in vivo*, but with remarkable brain-region-dependent effects on mitochondrial dynamics, spontaneous Ca^2+^ fluxes and astrocytic and microglial reactivity. Thus, AAV-Mito-PstI is a novel tool to explore the relationship between astrocytic mitochondria network dynamics and astrocytic mitochondria Ca^2+^ signaling in a brain-region-selective manner.

## RESULTS

### Mito-PstI targeted to astrocytes depletes and deletes mtDNA *in vivo*

Prior work showed that targeting of PstI to mitochondria in murine cells led to mtDNA depletion and a deletion of mtDNA between the two PstI restriction sites that are specifically present in murine mtDNA (Fukui & Moraes, 2009; Pickrell et al., 2011; Srivastava & Moraes, 2005). Based on this, we designed an AAV in which the well-established, GfaABC1D astrocyte-specific promoter (Y. Lee, Messing, Su, & Brenner, 2008) drives expression of PstI, specifically targeted to astrocytic mitochondria via a mitochondria signal sequence. In addition, we utilized previously published AAV reporter constructs to specifically drive expression of either GCaMP6f (Mito-GcaMP6f) or GFP (Mito-GFP) in astrocytic mitochondria (Fig. 1A) (Huntington & Srinivasan, 2021). The expression pattern of Mito-GFP in the DLS and the HPC dentate gyrus (Fig 1B) demonstrated the highly selective astrocyte specificity allowed by these vectors. As a first step, we determined whether our Mito-PstI construct would lead to depletion of mtDNA in astrocytes *in vivo*. To do this, we introduced 1 x 10^10^ gc of our AAV via stereotaxic surgery unilaterally into the DLS of 2-to 3-month-old mice, and after ∼3 wk, microdissected the injected DLS and isolated total DNA from this tissue and from DLS of age-matched, uninjected mice that served as control. Mitochondrial DNA content in the samples were quantified as the mtDNA/nuclear DNA ratio by sybr green quantitative RT-PCR (Venegas, Wang, Dimmock, & Wong, 2011) using primers to Cytochrome b (*Mt-cyb*, white arrows in Fig. 1C), a mitochondrial gene outside the PstI sites, and Erk2 (*Mapk1*), as nuclear gene. As seen in Figure 1D, Mito-PstI-infected tissue had on average 60.1 ± 4.6 % mtDNA content relative to the control (n=3; p=0.0044, t-test). ∼40% depletion in mtDNA content *in vivo* is consistent with selective targeting of astrocytic mtDNA, and not of mtDNA of all cell types in the DLS. Since mitochondria-targeted PstI has also been shown to induce mtDNA deletions by recombination in neurons and muscle (Pickrell et al., 2011; Srivastava & Moraes, 2005), we used two pairs of nested primers flanking the two PstI sites in mitochondria (black and gray arrows in Fig. 1C) to amplify the region between them by standard PCR. While the expected 4512 bp fragment from wildtype mtDNA is too large to be detected with our amplification conditions (see Methods), the much smaller DNA bands generated by recombination following PstI digestion would be amplified using our protocol. In agreement with this rationale, we detected a ∼700 bp fragment only in samples from mice treated with Mito-PstI *in vivo* (Fig. 1E). The size of this fragment was similar to that reported by the Moraes group using primers that also flanked the PstI sites in experiments in which PstI was expressed in dopaminergic neurons (Pickrell et al., 2011). The bands were excised, extracted, and sent for sequencing, which confirmed that recombination indeed took place between the two mtDNA PstI sites (Supplementary Material Figure 1). Thus, Mito-PstI targeted to astrocytes *in vivo* causes significant depletion of mtDNA and recombination events consistent with prior results in neurons (Pickrell et al., 2011).

**Figure 1.**
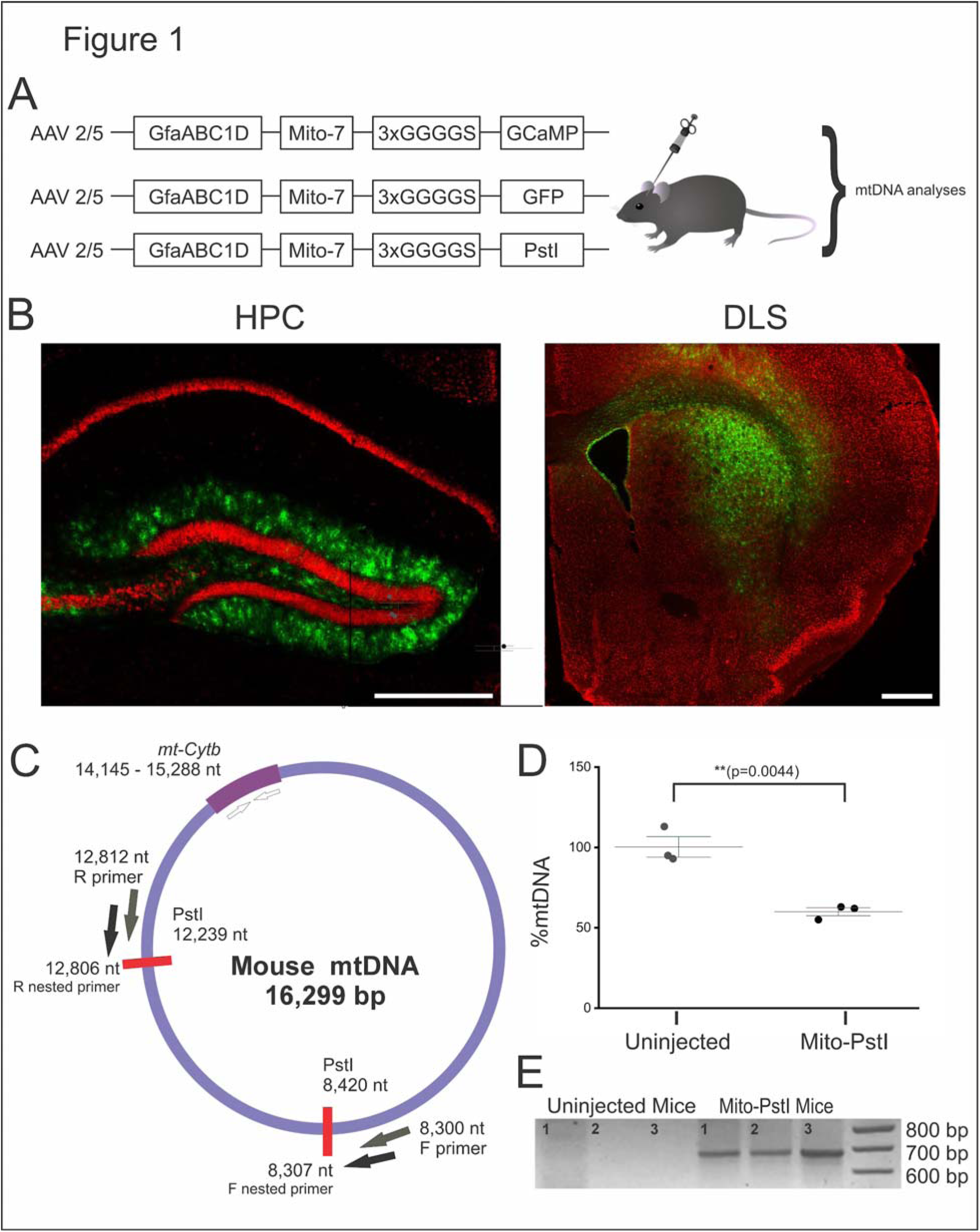
Mito-PstI depletes and deletes astrocytic mtDNA *in vivo*. (A) Schematic showing the AAV 2/5 vectors for stereotaxic injection into specific brain regions. GfaABC1D: *Gfap* promoter. Mito-7: COX8A mitochondrial signal sequence. 3XGGS: 12 amino acid linker. **(B)** Images depicting Mito-GFP fluorescence on NeuN stained HPC and DLS tissue, respectively. GFP expression was highly astrocyte specific. Scale bar = 500 μm. **(C)** Diagram for mtDNA genome with location of PstI sites (red bars), PCR primers for *Cytochrome b* (white arrows), and for detection of possible deletions (black and gray arrows). Nucleotide (nt) positions are indicated. **(D)** Relative mtDNA content quantification showing a ∼40% reduction in mtDNA content in Mito-PstI-treated tissue. Each data point is midbrain homogenate from a single, male mouse. Errors bars areL±LSEM, p-value for the relative mtDNA content quantification is based on a two-sample t-test, n = 3 male mice per condition. **(E)** Detection of potential deletions. Size markers in the right-most lane; ∼700 bp fragment, suggestive of mtDNA recombination, only observed in Mito-PstI treated samples (n=3/condition).

### Mito-PstI causes significant morphological changes to astrocytic mitochondria in DLS, but not in HPC

We recently demonstrated robust morphological and functional differences in astrocytic mitochondria between the hippocampus and striatum (Huntington & Srinivasan, 2021). Therefore, in this study, we sought to utilize AAV-Mito-Pst1 as a tool to determine if damaging astrocytic mtDNA within the HPC and DLS would result in divergent effects on astrocytic mitochondria in these two brain regions. Thus, we unilaterally co-injected a 1:1 mixture of AAV-Mito-PstI and AAV-Mito-GFP (1 x 10^10^ gc each AAV, 2 x 10^10^ gc total) either into the DLS or the dentate gyrus of the hippocampus (HPC) of 2–3-month-old mice. As a control, we injected mice with 2 x 10^10^ vg of AAV-Mito-GFP in either DLS or HPC. This AAV vector utilized the same GfaABC1D promoter and mitochondrial signal sequence that was used to target PstI (Fig 1A), which allowed clear visualization of astrocytic mitochondria (Fig 2A-D). After ∼3 wk, brains were dissected, sectioned in a cryostat and these were imaged in a confocal microscope. Images of individual Mito-GFP-expressing astrocytes were taken (Fig. 2A, 2B, 2C, 2D) and analyzed using ImageJ. We first quantified the number, size, and combined area of mitochondrial particles in individual astrocytes within each of the two brain regions (Fig. 2E, 2F, 2G). Results revealed that baseline mitochondrial number and size were significantly different between DLS and HPC as Mito-GFP control mitochondrial particles in the DLS were ∼2-fold more abundant (Fig 2E) and ∼5-fold larger (Fig 2F) than those in HPC (Table 1). Under control Mito-GFP injected conditions, no significant differences were observed in whole network area of mitochondria between the HPC and DLS (Fig. 2G, Table 1). Astrocytic PstI expression did not significantly alter the number, size and combined area of mitochondrial particles in the HPC, however, by contrast, we observed significant changes in the DLS. Thus, in the DLS, Mito-PstI-treated astrocytes had more mitochondria (Fig. 2E, Mito-GFP+PstI: 849 ± 49; Mito-GFP: 540 ± 23; p<0.0001, Mann Whitney test), that were smaller (Fig. 2F, Mito-GFP+PstI: 4.4 ± 0.39 µm^2^; Mito-GFP: 8.5 ± 0.83 µm^2^; p<0.0001, Mann Whitney test) than mito-GFP expressing control astrocytes. Additionally, Mito-PstI-expressing astrocytes in the DLS showed a significantly larger area of mitochondrial coverage per astrocyte (609 ± 69 µm^2^) than the Mito-GFP control group (331 ± 40 µm^2^; p=8.46E-4, Mann Whitney test). Thus, these baseline morphological parameters suggest that astrocytic mitochondria in DLS and HPC show heterogeneity at baseline in mitochondrial morphology, as well as diverging structural changes in mitochondria following astrocytic mtDNA damage. Of note, DLS mitochondria, in particular, appeared more fragmented with Mito-PstI expression.

**Figure 2.**
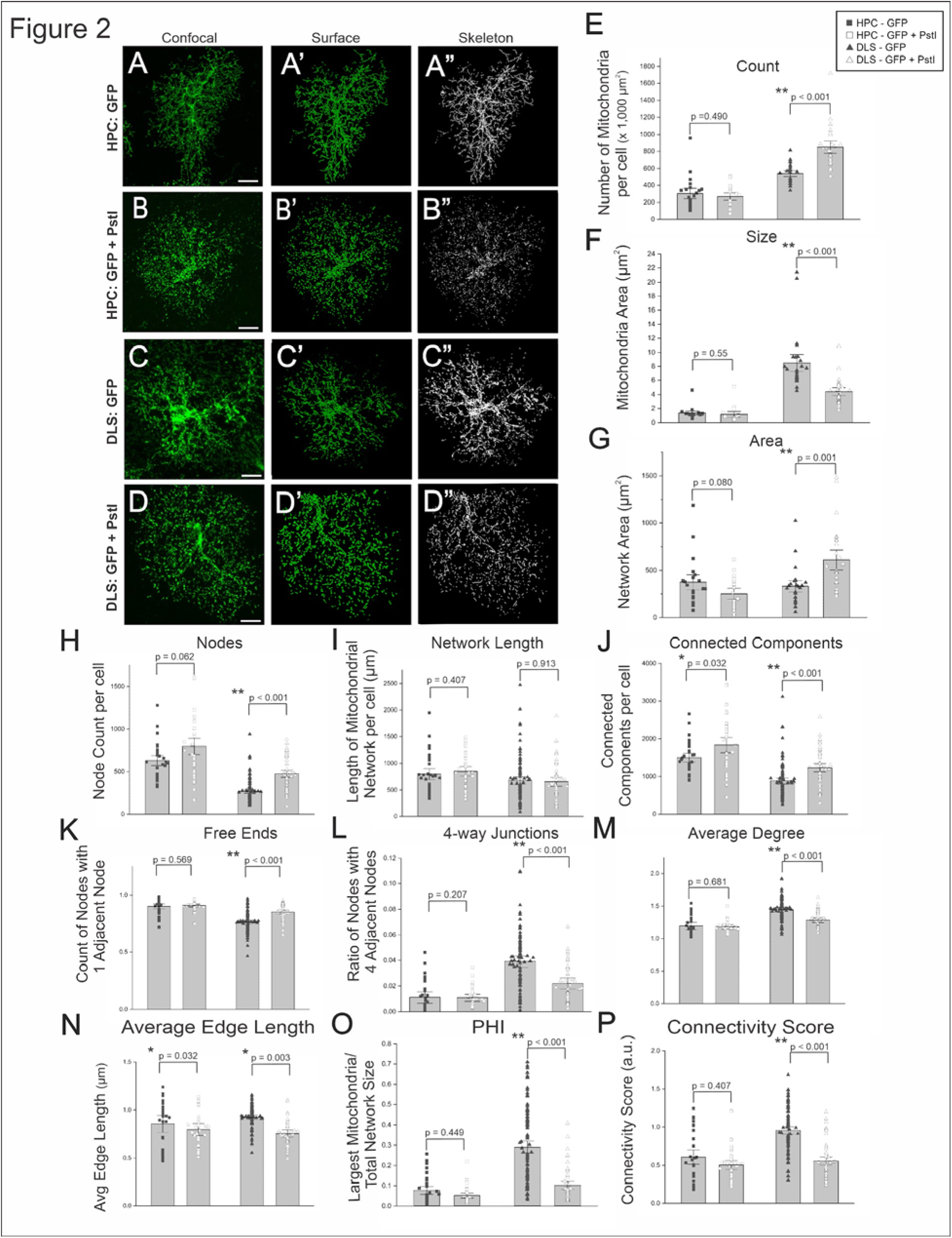
Mito-PstI causes significant morphological changes to astrocytic mitochondria in DLS, but not in HPC. Representative confocal images **(A-D)**, MitoGraph surface reconstructions **(A’-D’)**, and MitoGraph mitochondrial network skeletons (A”-D”) of Mito-GFP expression in HPC (A-B) and DLS (C-D) astrocytic mitochondria. Scale bars = 10 μm. Mitochondrial particle count **(E)**, size **(F)**, and network area **(G)** as quantified using ImageJ. Mitochondrial network connectivity parameters assessed using MitoGraph **(H-P)**. For image analyses, each data point is a single astrocyte. Data expressed as mean ±LSEM, p-values are based on a two-sample t-test or Mann Whitney test, depending on normality. For (E-G), n = 21 Mito-GFP only astrocytes and 17 Mito-GFP + PstI astrocytes in the HPC, n = 25 astrocytes per condition in the DLS. For (H-P), n = 30 astrocytes per condition in the HPC, n = 93 Mito-GFP-only astrocytes and n = 43 Mito-GFP + Mito-PstI astrocytes in the DLS. Brain slices were taken from 3-5 mice, using both males and females, per condition per brain region.

**Table 1.** Baseline comparison of GFP labeled mitochondrial count, size, and area in the HPC and DLS. Average values +/− SEM are represented for each of these parameters.

|  | Mito-GFP Only |  |  |
| --- | --- | --- | --- |
|  | HPC | DLS | P-value |
| Count<sup>#</sup> | 302 +/- 41 | 540 +/- 23 | <0.0001 |
| Size<sup>\$</sup> | 1.3 +/- 0.2 | 8.5 +/- 0.8 | <0.0001 |
| Area<sup>#</sup> | 374 +/- 53 | 330 +/- 40 | 0.6776 |

\*All p-values are based on a Mann-Whitney test.
#Counts are derived as the number of mitochondria per astrocyte

To further study the morphological effects of Mito-PstI on astrocytic mitochondria, we next used MitoGraph, a software package specifically designed to quantitatively analyze mitochondrial networks (Harwig et al., 2018). The confocal images were processed using MitoGraph to reconstruct the mitochondrial network surfaces (Fig. 2A’-D’) and skeletons (Fig. 2A”-D”). A number of network connectivity parameters (Fig. 2H-P) resulted from the skeletonizing of the segmented mitochondrial network into a set of edges (i.e., mitochondrial tubules), and nodes (i.e., endpoints and branch points). Edges are the connection between nodes and individual mitochondrial particles are referred to as connected components. In the HPC, node (Fig. 2H, Mito-GFP: 1504.8 ± 78.87, Mito-GFP+PstI: 1835 ± 133.1, p=0.06173, Mann Whitney test) and connected component counts (Fig. 2J, Mito-GFP: 632 ± 40, Mito-GFP+PstI: 797 ± 63, p=0.03107, t-test) were elevated in the Mito-GFP + Mito-PstI group compared to Mito-GFP controls, however, the overall connectivity scores were not different from the controls (Fig. 2P). This suggests that Mito-PstI expression failed to significantly change morphological characteristics of the mitochondrial network within astrocytes in the HPC. In stark contrast, the mitochondrial network in DLS astrocytes for the Mito-GFP + Mito-PstI group had a significantly higher number of nodes (Fig. 2H, Mito-GFP: 883 ± 49, Mito-GFP+PstI: 1234 ± 74; p<0.0001, Mann Whitney test), connected components (Fig. 2J, Mito-GFP: 270 ± 15, Mito-GFP+PstI: 474 ± 29; p<0.0001, Mann Whitney test) and free ends (Fig. 2K, Mito-GFP: 0.76 ± 0.01, Mito-GFP+PstI: 0.85 ± 0.012; p<0.0001, Mann Whitney test) than control. In addition, Mito-Pst1 expressing DLS astrocytes had significantly fewer 4-way junctions (Fig. 2L, Mito-GFP: 0.0391 ± 0.0022, Mito-PstI: 0.022 ± 0.0023; p<0.0001, Mann Whitney test), lower Average Degree (Fig. 2M, Mito-GFP: 1.44 ± 0.018, Mito-GFP+PstI: 1.28 ± 0.020, p<0.0001, Mann Whitney test), Average Edge Length (Fig. 2N, Mito-GFP: 0.925 ± 0.0117 µm, Mito-GFP+PstI: 0.754 ± 0.0195 µm, p<0.0001, Mann Whitney test), PHI (Fig. 2O, Mito-GFP: 0.29 ± 0.02, Mito-GFP+PstI: 0.102 ± 0.013; p<0.0001, Mann Whitney test) and connectivity score (Fig. 2P, Mito-GFP: 0.96 ± 0.033, Mito-GFP+PstI: 0.56 ± 0.036; p<0.0001, Mann Whitney test). Importantly, the connectivity score is a single measure, intended to capture the diverse nature of the mitochondrial network, calculated by using as numerator the sum of parameters elevated in highly fused networks (e.g., PHI: relative size of the largest connected mitochondrial component compared to the total mitochondrial size, average edge length, and average degree), and as denominator the sum of parameters elevated in highly fragmented networks (e.g., nodes, edges, and connected components). A reduction in connectivity score tightly correlates with increased mitochondria fission (Harwig et al., 2018). Thus, these data suggest that Mito-PstI specifically causes increased fission, or decreased fusion, of mitochondrial networks within astrocytes in the DLS.

### Mito-PstI causes a striking divergence in the kinetics of spontaneous mitochondrial Ca^2+^ influx events between astrocytes in the HPC versus DLS

We previously reported that astrocytic mitochondria display robust, spontaneous Ca^2+^ influx events in situ with baseline inter-regional differences between astrocytes in the DLS and HPC (Huntington & Srinivasan, 2021). Thus, to evaluate the effects of PstI on spontaneous mitochondrial Ca^2+^ events, we unilaterally co-injected the DLS or the HPC dentate gyrus of 2–3-month-old mice with a mixture of AAV-Mito-PstI (1 x 10^10^ gc) and AAV-Mito-GCaMP6f (2 x 10^9^ gc). As control, we injected other mice with 2 x 10^9^ vg of AAV-Mito-GCaMP6f in either DLS or HPC. This virus delivered the GCaMP6f Ca^2+^ indicator (Helassa, Podor, Fine, & Török, 2016) selectively to astrocytic mitochondria in the two brain regions, (Fig 1A; (Huntington & Srinivasan, 2021)). After 3-4 wk, Ca^2+^ imaging of individual astrocytes was performed in live brain slices as detailed in Methods, and recordings were analyzed with the AQuA software package (Wang et al., 2019). In preliminary analysis of the data, we discovered we could bin the observations into two distinct types of Ca^2+^ influx events, based on their propagation distance. Considering the size of mitochondria (∼1 μm in length), and movement in the x-y axis due to mitochondrial and optical jitter, we defined non-propagating events as those with a traveling distance of 0.99 μm or below, while propagating events were those with a traveling distance of 1 μm or higher. Figure 3, and Supplementary Movies 1-4, show representative examples of these types of events.

**Figure 3.**
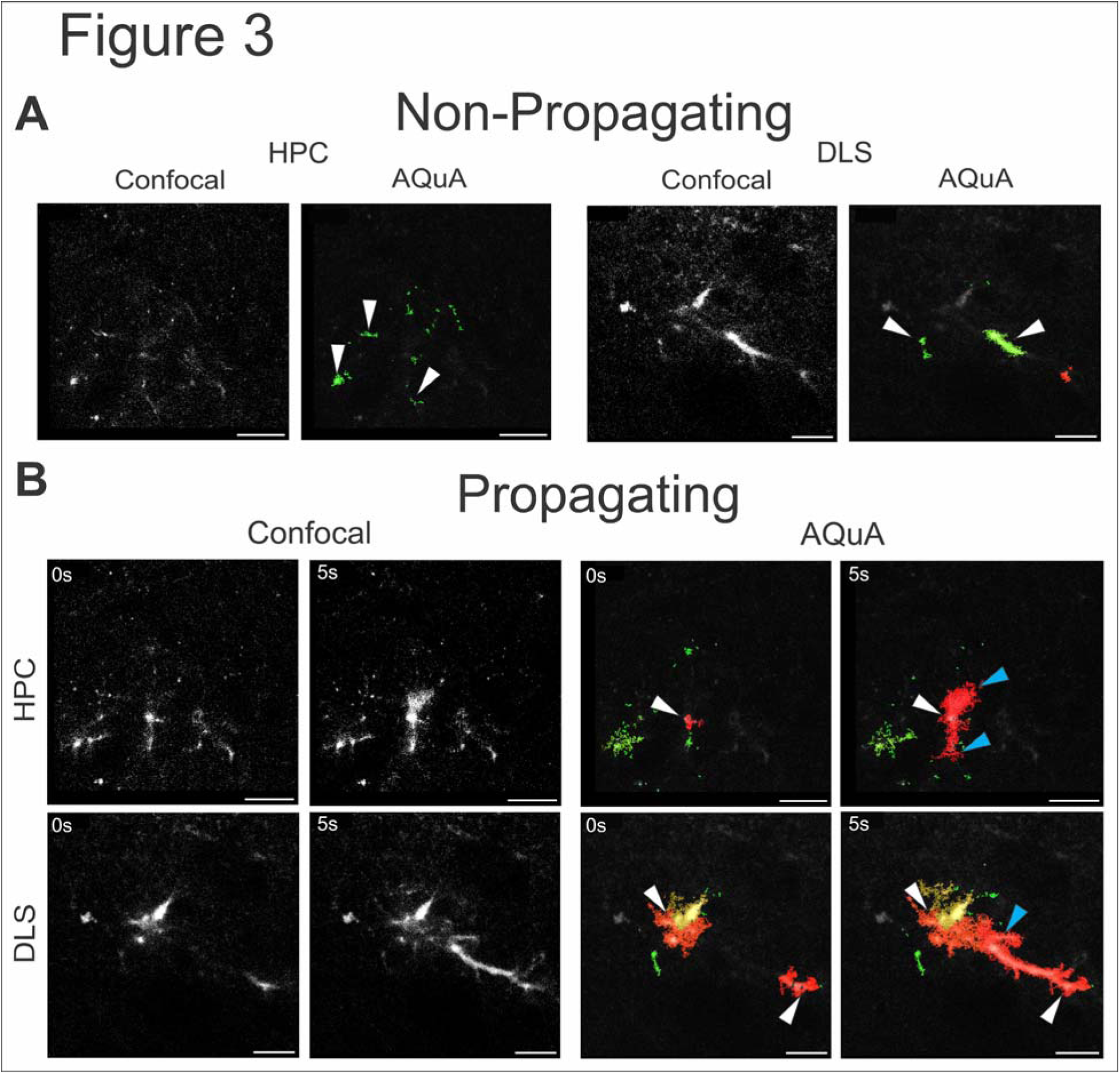
Representative images of Ca^2+^ influx events separated by type. (A) Confocal (left) and AQuA (right) representation of Non-Propagating astrocytic mitochondrial Ca^2+^ influx in live HPC (left) and DLS (right) brain slices. **(B)** Confocal (left) and AQuA (right) representation of Propagating astrocytic mitochondrial Ca^2+^ influx with a 5 second interval in the HPC (top) and DLS (bottom) live brain slices. White arrows depict initiation of events, while blue arrows indicate ending-point of propagating events. Propagating events are color-coded red, whereas Non-Propagating events are color-coded green by AQuA. Scale bars = 20 μm.

Using these parameters to discriminate non-propagating from propagating events, we examined whether the introduction of Mito-PstI would cause changes in mitochondrial Ca^2+^ influx kinetics dependent on this segregation. Overall, we found that non-propagating events vastly outnumbered propagating events by about 10:1 in frequency, regardless of the brain region examined (Fig 4). At baseline, we observed significant differences in kinetic parameters for non-propagating and propagating events between the HPC and DLS. Specifically for non-propagating events, the DLS displayed a significant ∼20% greater amplitude in mitochondrial calcium events, while duration of events in the DLS were ∼3-fold greater than the HPC (Table 2). We also observed significant baseline changes between the HPC and DLS for propagating events. Specifically, the DLS showed ∼4-fold greater duration of events, along with ∼40% reduction in propagation distance (Table 2). As for PstI effects, non-propagating events in the DLS showed a significant increase in frequency (Mito-GCaMP: 147.70 ± 36.481, Mito-GCaMP+PstI: 336.27 ± 278.53, p=0.012, t-test), while amplitude (Mito-GCaMP: 1.23 ± 0.044, Mito-GCaMP+PstI: 1.11 ± 0.036, p=0.036) and duration (Mito-GCaMP: 2.88 ± 0.104, Mito-GCaMP+PstI: 2.608 ± 0.083, p = 0.048, t-test) were reduced. By contrast, in the HPC (Fig 4A), non-propagating event frequency and duration were unaffected by PstI, while amplitude was significantly increased (Mito-GCaMP: 0.997± 0.070, Mito-GCaMP+PstI: 1.188 ± 0.062, p=0.037, Mann-Whitney test).

**Figure 4.**
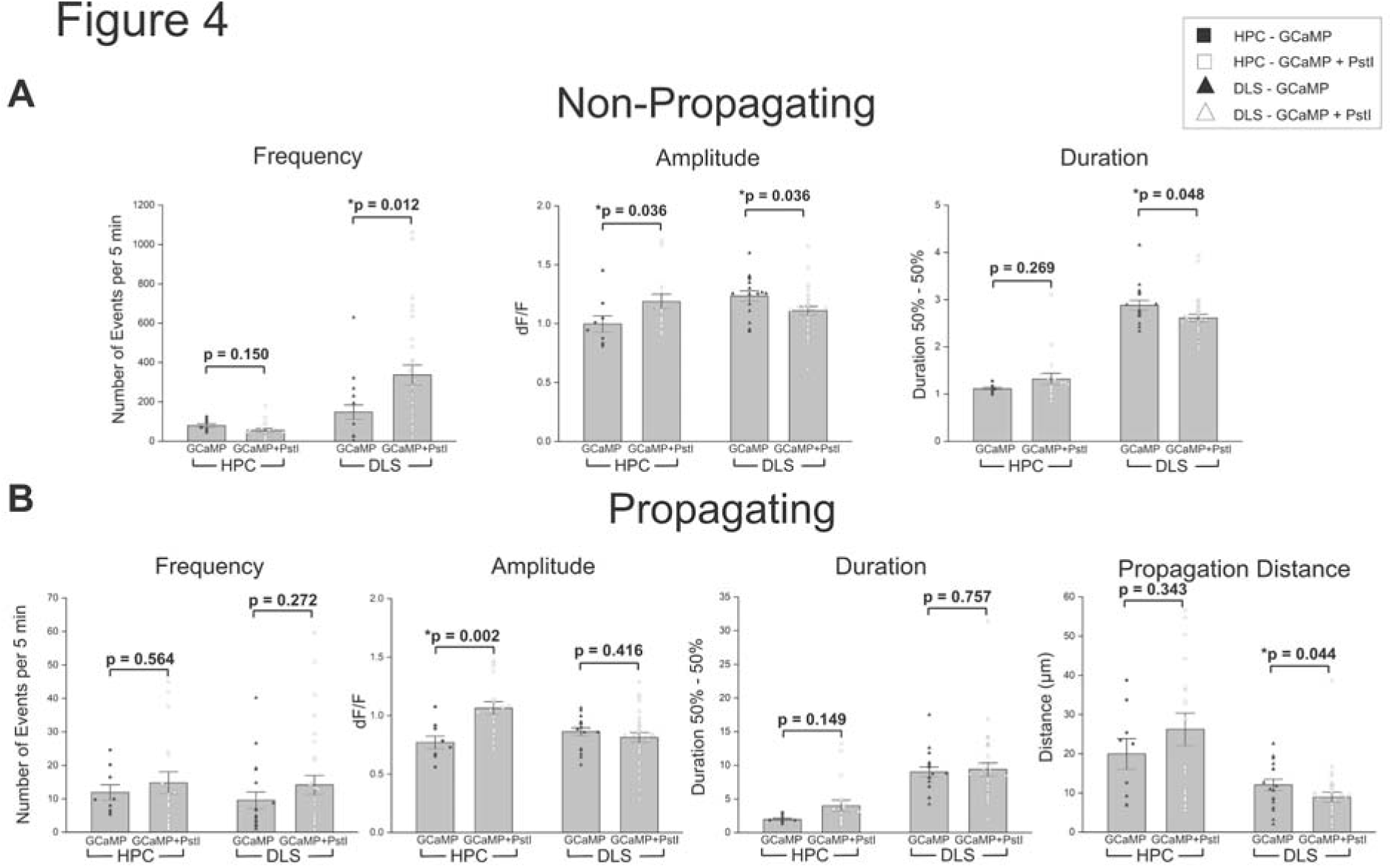
Mito-PstI causes a striking divergence in the kinetics of spontaneous mitochondrial Ca^2+^ influx events between astrocytes in the HPC versus DLS. Graphs depicting differences in frequency, amplitude, duration and propagation distance between Mito-GCaMP and Mito-GcaMP+PstI conditions for the two types of Ca2+ events: Non-propagating **(A)** and Propagating **(B)**. Ca^2+^ influx from live HPC and DLS brain slices was analyzed utilizing the AQuA software plug-in for ImageJ. For image analyses, each data point is a single astrocyte. Data expressed as mean ±LSEM, p-values are based on two-sample t-test or Mann-Whitney test. For HPC, n = 3 mice, 9 cells (Mito-GCaMP) and n = 10 mice, 18 cells (Mito-GCaMP+PstI). For DLS, n = 7 mice, 17 cells (Mito-GCaMP) and n = 8 mice, 30 cells (Mito-GCaMP+PstI)

**Table 2.** Baseline comparison of frequency, amplitude, and duration of Mito-GcaMP6f-based fluorescent non-propagating and propagating Ca^2+^ influx signals in HPC and DLS astrocytic mitochondria. Comparisons of propagation distance in the HPC and DLS are also shown. Average values +/− SEM are represented for each of these parameters.

|  |  | Mito-GCaMP6f Only |  |  |
| --- | --- | --- | --- | --- |
|  |  | HPC | DLS | P-value |
| Non-Propagating<sup>#</sup> | Frequency | 79.12 +/- 9.9 | 147.7 +/- 36 | 0.777 |
|  | Amplitude | 0.997 +/- 0.07 | 1.2 +/- 0.04 | 0.006 |
|  | Duration | 1.1 +/- 0.03 | 2.9 +/- 0.10 | <0.0001 |
| Propagating<sup>#</sup> | Frequency | 11.9 +/- 2.3 | 9.6 +/- 2.5 | 0.089 |
|  | Amplitude | 0.77 +/- 0.05 | 0.86 +/- 0.03 | 0.140 |
|  | Duration | 1.9 +/- 0.57 | 9 +/- 3.1 | <0.0001 |
|  | Distance | 20 +/- 11.7 | 12.1 +/- 6.05 | 0.028 |

\*p-values are based on either a Mann-Whitney or two-sample t-test.
#Units for frequency, events per 5 min.; amplitude, dF/F; duration, s; distance, $\mu\text{m}$ .

For propagating events, we found that propagation distance of mitochondrial Ca^2+^ influx events significantly decreased only in the DLS (Mito-GCaMP: 12.094 ± 1.427 µm, Mito-GCaMP+PstI: 8.983 ± 1.284 µm, p=0.044, Mann Whitney test), but remained unchanged in the HPC. Although propagating event frequency and duration remained unchanged in both brain regions after PstI treatment (Fig 4B), the amplitude of propagating events showed a small but significant increase only in the HPC of PstI-treated cells (Mito-GCaMP: 0.770 ± 0.054, Mito-GCaMP+PstI: 1.06 ± 0.055, p=0.002, t-test). Thus, following mtDNA damage induced by mitochondrial PstI expression, the kinetics of spontaneous mitochondrial Ca^2+^ influx events in astrocytes showed striking brain region-specific differences between the HPC and DLS. Importantly, significant differences were observed for non-propagating events in the DLS but not in the HPC.

### Mito-PstI causes astrocytes to become reactive in the HPC and DLS

Astrocytes adopt a reactive state, known as astrogliosis, in response to infection, injury or chronic disease, which involve variable morphological, functional, and molecular changes (Escartin et al., 2021). Since astrocytic mtDNA depletion has been shown to trigger astrogliosis (Ignatenko et al., 2018), and glial fibrillary acidic protein (GFAP) gene and protein induction are classical markers of astrogliosis (Yang & Wang, 2015), GFAP content was assessed via immunostaining in both the HPC and DLS (Fig. 5A-D). In the HPC, both GFAP staining area (Fig. 5E, Mito-GFP: 319 ± 28 µm^2^, Mito-GFP+PstI: 523 ± 72 µm^2^, p=0.0137, Mann Whitney test) and intensity (Fig. 5F, Mito-GFP: 295 ± 11, Mito-GFP+PstI: 384 ± 29, p=0.00747, t-test) increased in the Mito-GFP + Mito-PstI mice. The staining area also increased in the PstI-expressing DLS samples (Fig. 5E, Mito-GFP: 607 ± 43 µm^2^, Mito-GFP+PstI: 770 ± 46 µm^2^, p=0.0122, t-test), however, the intensity decreased (Fig. 5F, Mito-GFP: 330 ± 86, Mito-GFP+PstI: 265 ± 82, p=0.00832, t-test). Thus, overall these data suggest that astrocytes in both HPC and DLS become reactive in response to Mito-PstI expression.

**Figure 5.**
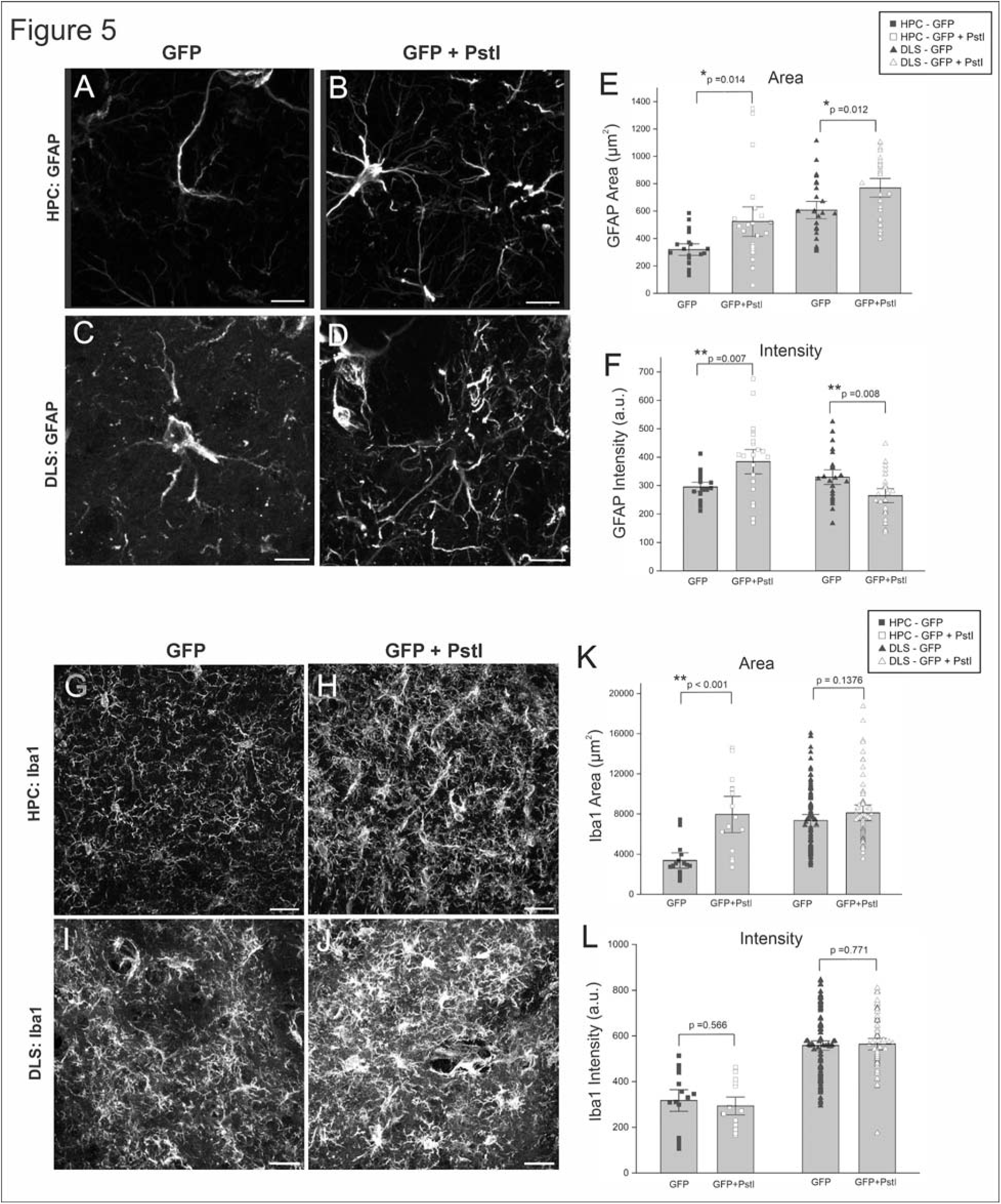
The HPC and DLS display heterogenous differences in astrocytic and microglial reactivity to Mito-PstI. Representative confocal images of GFAP staining in HPC **(A-B)** and DLS **(C-D)** astrocytes expressing either Mito-GFP (A, C) or Mito-GFP+PstI (B, D). ImageJ quantification of GFAP area **(E)** and intensity **(F)** per field of view (FOV). Scale bars = 10 μm. Representative confocal images of Iba1 staining in HPC **(G-H)** and DLS **(I-J)** sections expressing either Mito-GFP (G,I) or Mito-GFP+PstI (H,J). ImageJ quantification of Iba1 area **(K)** and intensity **(L)**. Each data point is a single FOV taken from 1-4 brain slices from 3-5 mice, male and female, per condition per brain region (HPC and DLS). Data expressed as mean ±LSEM, p-values are based on a two-sample t-test or Mann Whitney, based on normality. For GFAP, n = 29 FOV per condition in the HPC, 25 FOV per condition in the DLS. For Iba1, n = 18 FOV per condition in the HPC, 27 FOV per condition in the DLS.

### Mito-PstI elicits differential microglial activation in the HPC and DLS

Microglia, the resident immune cells of the CNS, respond to perturbations of their surroundings, such as injury or neurodegenerative disease, by increasing their numbers and undergoing a series of morphological changes. This response, known as microgliosis, is very dynamic and heterogeneous, depending on multiple variables such as: brain region, age, sex, and specific pathological agent (Paolicelli et al., 2022). Chronic loss of astrocytic mtDNA triggers astrogliosis, and secondarily, microgliosis (Ignatenko et al., 2018). Therefore, we asked whether our astrocyte-targeted Mito-PstI tool would elicit microglia reactivity. We performed immunostaining for the microglial marker ionized calcium-binding adapter molecule 1 (Iba1) in both HPC and DLS slices, which received either a Mito-GFP or a Mito-GFP + Mito-PstI AAV injection (Fig. 5 G-J). Quantification showed that Iba1 staining area significantly increased only in the PstI-treated HPC (Fig. 5K, Mito-GFP: 3370 ± 391 µm^2^, Mito-GFP+PstI: 7952 ± 918 µm^2^, p=1.06E-4, Mann Whitney test), but not in the PstI-treated DLS, while Iba1 intensity (Fig. 5L) remained unchanged following PstI treatment in both brain regions. This may indicate increase recruitment of microglia to the HPC due to Mito-PstI expression in astrocytes, without alterations of Iba1 expression per cell. Thus, together, these data suggest that targeting PstI to astrocytic mitochondria triggers microgliosis in the HPC, but perhaps not in the DLS.

## DISCUSSION

The data show that PstI targeted to astrocytic mitochondria induces mtDNA loss and recombination *in vivo.* This has brain-region specific effects on astrocytic mitochondrial network dynamics and spontaneous Ca^2+^ influx events, with accompanying astrocytic activation and microglia reactivity, also brain-region dependent.

AAV-Mito-GFP, AAV-Mito-GCaMP6f and AAV-Mito-PstI all used the same genome/capsid pairing (AAV2/5), GfaABC1D promoter, and mitochondrial signal sequence to deliver their respective cargoes to astrocytic mitochondria (Fig 1A). When used alone, AAV-Mito-GFP clearly labelled astrocytic mitochondria (Fig 1B, Fig 2A, 2C) without targeting mitochondria in any other cell, so was demonstrated previously for AAV-Mito-GCaMP (Huntington & Srinivasan, 2021). Thus, there is little doubt that AAV-Mito-PstI selectively delivered this restriction enzyme to just astrocytic mitochondria either when used alone (Fig 1), or in combination either with AAV-Mito-GFP (Fig 2) or AAV-Mito-GCaMP (Fig 3 and 4). The degree and nature of mtDNA depletion and deletion yielded by the AAV tool used here to deliver PstI to astrocytes, are remarkably similar to those reported by Moraes and colleagues, who delivered PstI to neurons (Fukui & Moraes, 2009; Pickrell et al., 2011) and myofibers (Srivastava & Moraes, 2005), using tissue-specific transgenic approaches. This suggests that the mechanisms that lead to PstI-induced mtDNA loss and recombination, which likely involve DNA double-strand breaks, exonuclease activity and homologous and non-homologous DNA repair systems (Rong et al., 2021), are conserved across these cell types. A different approach to generate mtDNA loss in astrocytes was the conditional deletion of *Twinkle*, a nuclear encoded gene necessary for mitochondrial genome replication (Ignatenko et al., 2018). In this case, mtDNA depletion was achieved by defective replication. Interestingly, this progressive astrocytic mtDNA attrition led to astrogliosis, secondary microgliosis and neuronal loss. AAV-Mito-PstI treatment targeted to astrocytes also led to astrocytic activation and microgliosis, particularly in the HPC. We have yet to look at neuronal loss or other neuronal-specific effects with our tool. An advantage of the AAV-Mito-PstI tool over the conditional gene targeting approach used by Ignatenko and colleagues, is that it allows for brain-region-specific targeting of astrocytic mtDNA, which we have exploited here to show differential effects on mitochondrial network morphology and spontaneous Ca^2+^ influx in the DLS and HPC.

Simple quantification of mitochondrial particle number (Fig 2E) and size (Fig 2F), which respectively showed an elevation and a diminution in Mito-GFP+PstI samples relative to Mito-GFP control, strongly suggested PstI-driven augmentation of mitochondrial fragmentation, consistent with elevated mitochondrial fission (or reduced fusion) in DLS, but not in HPC. This was confirmed by the more sophisticated analysis with Mitograph, which yielded a significant decrease in the DLS connectivity score, in particular (Fig 2P), which has been experimentally linked to increased mitochondria fission (Harwig et al., 2018). In all cells (Youle & van der Bliek, 2012), in general, and astrocytes (Stephen et al., 2014), in particular, mitochondria fission, fusion and mitophagy form a tripartite quality control system that ensures eventual removal of damaged, non-functional mitochondria from the mitochondrial network of a cell. These processes are triggered by many cellular stresses including those that damage mtDNA like oxidative stress, and in our case, double-strand DNA breaks (Youle & van der Bliek, 2012) (Rong et al., 2021). It is intriguing that astrocytes in DLS and HPC differ so significantly regarding this homeostatic response. In this context, consistent with our previous results in the HPC CA1 region (Huntington & Srinivasan, 2021), control mitochondria in the astrocytes of the HPC dentate gyrus appeared smaller and less abundant than in the DLS (Fig 2 E,F; Table 1). It is tempting to suggest that these baseline morphological properties may account, in part, for the relative resilience of the HPC mitochondrial network to fission by PstI treatment. Interestingly, the connectivity index of the DLS astrocytic mitochondrial network was brought down to about the same value as that for the HPC after PstI treatment (Fig 2P).

Our prior study comparing DLS and CA1 HPC astrocytes found interregional baseline differences in spontaneous Ca^2+^ influx events (Huntington & Srinivasan, 2021). Even though, AQuA was not used for the analysis of the data in that report, it is likely that given the 10:1 predominance of Non-Propagating events reported here, most of the events analyzed in that previous study were Non-Propagating as well. Event frequency in CA1 HPC astrocytic mitochondria was about half of that in the DLS, which matches really well with what we found in the dentate gyrus here (Fig 4A). The analysis here also revealed important baseline differences particularly in event duration, ∼4 times as large in the DLS relative to the HPC, and in propagation distance, ∼2 times shorter in the DLS (Fig 4A,4B; Table 2).

Lower mitochondrial abundance in the HPC relative to DLS (Fig 2E; Table 1) might account for decreased baseline frequency of Non-Propagating Ca^2+^ events in the former relative to the latter (Fig 4A). Likewise, longer event duration in the DLS might be related to increased baseline connectivity in its mitochondrial network. Increased mitochondria fission in PstI-treated DLS may also account for the decrease in Non-Propagating event duration and the increase in Non-Propagating event frequency (Fig 4A), and perhaps, for the reduction of propagation distance in Propagating events (Fig 4B). It is difficult to relate the increase in event amplitude in HPC, and the decrease in Non-Propagating event amplitude in the DLS, to network connectivity status. Ca^2+^ influx in astrocytic mitochondria strongly depends on ER Ca^2+^ stores and not extracellular Ca^2+^ (Huntington & Srinivasan, 2021). It is likely that Ca^2+^ movement from ER to mitochondria in astrocytes involves, like in other cell types, mitochondria-associated ER membranes (MAMs) (S. Lee & Min, 2018). MAMs are present in astrocytes, where they have been shown structurally and functionally influenced by manipulation of mitochondrial dynamics (GLbel et al., 2020). Hence, it is reasonable to suggest that the disruption of the mitochondrial network induced by PstI, particularly in the DLS (Fig 2), is associated with alteration of MAMs, which in turn may underlie, in part, some of the more profound changes observed in spontaneous Ca^2+^ events in that brain region (Fig 4).

The induction of astrocyte reactivity by mtDNA loss using PstI (Fig 5 A-F) is not surprising given that conditional deletion of *Twinkle* in astrocytes, which also generate mtDNA depletion, did so as well (Ignatenko et al., 2018). Somewhat unexpected is that our data only support secondary microgliosis in the HPC and not the DLS (Fig 5 G-L). In addition to highlighting another brain region-specific difference for astrocytic mtDNA loss, this result raises the possibility of uncharacterized distinctions in reactivity to mtDNA damage between HPC and DLS astrocytes, which in turn may lead to different downstream responses by surrounding microglia. Diversity of astrocyte and microglia reactivity as a response to multiple kinds of insults, brain region, age, sex and other factors, is a well-established concept in the field (Escartin et al., 2021; Paolicelli et al., 2022).

In summary, the data demonstrate the utility of the novel AAV-Mito-PstI in inducing astrocytic mtDNA damage in a brain-region-specific manner, which produces stark interregional differences in mitochondrial network dynamics and spontaneous Ca^2+^ fluxes in astrocytes from DLS and HPC, two brain areas relevant to major neurodegenerative diseases, Parkinson’s and Alzheimer’s diseases. It remains to be investigated what other mitochondrial functions, i.e., ATP, ROS, lactate production, glutamate uptake, blood-brain barrier regulation, are affected by PstI-treatment and how their alteration could be mechanistically connected to the observed changes in mitochondrial dynamics, spontaneous Ca^2+^ fluxes, and astrocytic and microglial reactivity reported here, and to potential neuronal effects germane to Parkinson’s and Alzheimer’s diseases.

## MATERIALS AND METHODS

### Mice

Male and female C57BL/6 WT mice were obtained at 5-7 weeks old from Jackson Laboratories. Mice were housed on a 12 h light/dark cycle with *ad libitum* access to food and water. All animal experiments were conducted in accordance with Texas A&M University IACUC guidelines (Animal Use Protocol #2022-0252).

### Adeno-associated viral vectors (AAVs)

AAV 2/5 GfaABC1D-mito7-eGFP and AAV 2/5 GfaABC1D-mito7-GCaMP6f constructs were generated by Vector Builder (Chicago, IL) as described previously (Huntington & Srinivasan, 2021). A third AAV, AAV 2/5 GfaABC1D-mito7-PstI, was created in a similar manner, fusing the PstI coding sequence, optimized for mammalian codon use, to an N-terminal 87 bp Mito7 signal sequence from the mitochondrial COX8A subunit, driven by the astrocyte specific GfaABC1D promoter (Fig. 1A). We have verified genomes for all AAVs by sequencing. These vectors were designed to express each target protein only within astrocytic mitochondria.

### Stereotaxic surgery

Following prior protocols (Huntington & Srinivasan, 2021), stereotaxic surgeries were performed under continuous isoflurane (induction at 5%, maintenance at 1–2% vol/vol), administered via a gas syringe injection system (Kent Scientific). Glass injection pipettes (Fisher Scientific) were pulled using a Sutter P-2000 laser puller and tips were beveled using a Narishige EG-45 grinder.

For immunohistochemistry, 2×10^10^ genome copies (gc) of AAV were stereotaxically injected into the HPC or DLS. Control mice were injected with 2×10^10^ gc of AAV2/5 GfaABC1D-mito7-eGFP. Test animals were injected with 1×10^10^ gc of AAV2/5 GfaABC1D-mito7-eGFP and 1×10^10^ gc of AAV2/5 GfaABC1D-mito7-PstI.

For Ca^2+^ imaging, control animals were injected with 2×10^9^ gc of AAV2/5 GfaABC1D-mito7-GCaMP6f. Test animals were injected with 2×10^9^ gc of AAV2/5 GfaABC1D-mito7-GCaMP6f and 1×10^10^ gc of AAV2/5 GfaABC1D-mito7-PstI.

All injections were done using the Stoelting Quintessential Stereotaxic Injector (QSI). Mice were sacrificed 2-4 weeks post-injection for either perfusions or live-slice imaging. HPC coordinates were –2.0 mm posterior to bregma, +1.5 mm lateral to bregma, and –1.6 mm ventral to pial surface. DLS coordinates were +0.9 mm anterior to bregma, +1.8 mm lateral to bregma, and +2.5 mm ventral to the pial surface.

### mtDNA content analysis

DLS from two groups of three mice, which either had a stereotaxic injection of mito-PstI or were uninjected controls, were microdissected and pooled. DNA was extracted using the QIAGEN DNeasy Blood and Tissue Kit (QIAGEN; Germantown, MD; CAT# 69504). For quantitative PCR, Perfecta SYBR Green FastMix with ROX (QuantaBio; Beverly, MA; CAT# 95073) was used with Forward and Reverse Primers (Supplementary Material Table 1) for amplifying both Cytochrome b, a mtDNA gene, and Erk2, a nuclear DNA gene. The cycling parameters for amplification were as follows: 95°C for 7 min, [95°C for 10 sec, 60°C for 30 sec] 50 cycles, 95°C 15 sec, 60°C 1 min, 95°C 15 sec. Mitochondrial DNA content was calculated as described in (Venegas et al., 2011). Nested PCR was performed using SYBR Green FastMix (10 μL), Primers (Supplementary Material Table 1) were designed to sit outside the PstI cut sites (Fig 1C). Cycles were 95°C for 2 min, [94°C for 45 sec, 50°C for 45 sec, 60°C for 1 min] 35 cycles, 60°C for 7 min, 10°C hold. PCR product was run on a 1.5% agarose gel at 110V for 40 min. Bands were cut from the gel and sent to EtonBio (San Diego, CA) for sequencing.

### Immunostaining

Three-weeks after AAV injections, mice were anesthetized with isoflurane, transcardially perfused with 1X PBS, followed by 10% Formalin, and brains were extracted. Brains were post-fixed in 10% formalin at 4°C overnight before being moved to 30% sucrose solution for cryoprotection. Brains were set using Tissue Tek O.C.T. Compound (Sakura Finetek) and 40 μm sections were obtained using a cryostat (Epredia, CryoStar NX50).

For NeuN staining (Fig. 1B), following a 1X PBS wash, HPC and DLS sections were blocked and permeabilized in 10% normal goat serum (NGS, Abcam, ab7481) and 0.5% Triton X-100 (Sigma, cas# 9002-93-1) rocking gently for 45 min at room temperature. Sections were washed twice with PBS before incubation overnight at 4°C in primary mouse IgG2b anti-NeuN (1:500, Abcam cat# ab104224). Sections were washed twice the following day with 1X PBS followed by incubation in goat anti-IgG2b AlexaFlour 647 (1:1000, life technologies, ref# A21235). Slices were mounted using mounting media with DAPI (Fluoroshield, Abcam, ab104139).

The same protocol was used for GFAP and Iba1 staining (Fig. 5). Chicken anti-GFAP (1:2000, Abcam cat# ab4674) primary in 1% NGS + 0.05% Triton X-100 was used. For HPC slices, the secondary incubation was done in donkey anti-chicken AlexaFluor 647 (1:2000, JacksonImmuno cat#706-605-155) for one hour at room temperature. For DLS slices, goat anti-chicken AlexaFluor 594 (1:2000, Abcam cat# ab150176) was used.

For Iba1 staining in the HPC, rabbit anti-Iba1 (1:750, Wako cat#019-19741) was used as a primary antibody, and goat anti-rabbit AlexaFluor 594 (1:1500, Abcam cat#ab150080) was used for secondary. For DLS, rabbit anti-Iba1 (1:1000, Wako cat#019-19741) was used as a primary antibody, and goat anti-rabbit AlexaFluor 594 (1:2000, Abcam cat#ab150080) for secondary.

### Imaging

#### Imaging of Fixed Tissue

Fixed tissue sections of NeuN-stained HPC or DLS were imaged using an Olympus VS120 Virtual Slide Scanning System (Fig. 1B) equipped with a UPlanSApo 20X air objective. For Figures 2 and 5, fixed tissue sections of HPC and DLS were imaged using an Olympus FV3000 laser-scanning confocal microscope. 488, 561 and 647 nm LED-based excitation wavelengths were used for obtaining images. Images for MitoGraph analysis (Fig. 2) were obtained using a 60× oil immersion objective (N.A. 0.8), and 3× digital zoom. Z-stacks for these images consisted of 51 optical sections with a step size of 0.2 μm (Fig. 2). For HPC and DLS, GFAP and IBA1 images (Fig. 5) were obtained using a 60× oil immersion objective (N.A. 0.8) with 1x digital zoom. These images consisted of ∼25 optical sections with a step size of 0.5 μm. To enable quantification, confocal parameters such as excitation intensity, HV, gain and offset were maintained constant for each brain region.

#### Tissue preparation for live brain slicing

2 to 3-weeks following AAV injection, mice were decapitated and brains were extracted. 300 μm-thick coronal slices for HPC and DLS were obtained using a vibratome (Ted Pella Inc., D.S.K Microslicer ZERO 1 N). The slicing solution consisted of (in mM): 194 sucrose, 30 NaCl, 4.5 KCl, 1.2 NaH2PO4, 26 NaHCO3, 10 D-glucose, and 1 MgCl2 and saturated with 95% O2 and 5% CO2 (pH 7.2). All recordings of Ca^2+^ signals were performed in artificial cerebrospinal fluid (ACSF) made of (in mM): 126 NaCl, 2.5 KCl, 1.24 NaH2PO4, 26 NaHCO3, 10 D-glucose, 2.4 CaCl2, and 1.3 MgCl2 saturated with 95% O2 and 5% CO2 (pH 7.4).

#### Live imaging of acute mouse brain slices

An Olympus FV3000 upright laser-scanning confocal microscope equipped with a 40x water immersion objective (N.A. 0.45) was used for live slice imaging (Fig. 3). Confocal parameters (LED power, voltage, gain, offset, and aperture diameter) were maintained constant across all HPC or DLS GCaMP6f imaging sessions. All mitochondrial Ca^2+^ influx events were recorded at 1 frame per second for 5 minutes. During all live slice imaging sessions, temperature of the recording buffer and bath was maintained at 37°C using an in-line and bead bath heaters.

### Image analysis

Image analysis for Fig. 2E-G and Fig. 5 were performed using ImageJ (ver. 1.54). For Fig. 2E-G, a z projection of images was created using the max intensity function. The polygon tool was used to manually create a region of interest (ROI) around a single astrocyte. The area outside the cell was cleared and manually thresholded to encompass all GFP expressing mitochondrial particles, then smoothed and ROIs for mitochondrial particles were obtained from these images using the analyze particles function in ImageJ. ROIs obtained in this way were utilized for deriving particle count, size and area. For Fig. 5, ROIs were similarly obtained as described above, except that the images used were entire fields of view consisting of multiple cells. ROIs in Fig. 5 were utilized to obtain areas as well as intensity measures for GFAP and Iba1 staining.

### MitoGraph analysis

Single Mito-GFP expressing astrocytes within the HPC and DLS were analyzed using the MitoGraph package in ImageJ (Harwig et al., 2018). Maximum projections of single astrocytes were first obtained, and the astrocyte territory was manually delineated using the polygon tool. ROIs encompassing single astrocytes, obtained in this way were then applied to each of the 51 optical sections comprising the z-stack. Mito-GFP expressing mitochondria outside the ROI for a single astrocyte were then deleted in each of the 51 optical sections. For each astrocyte, z-stacks processed as described above were analyzed using the MitoGraph package run on the Texas A&M supercomputing cluster (Terra portal). MitoGraph outputs were analyzed using R statistical package. Mitochondrial network recreation files (skeleton, surface, and nodes) were visualized using the ParaView program, as shown in Fig. 2.

### AQuA Ca^2+^ imaging in acute mouse brain slices

The Astrocyte Quantitative Analysis (AQuA) (Wang et al., 2019) software plug-in for ImageJ was used to obtain and analyze Ca^2+^ events from Mito-GCaMP6f astrocytes. Ca^2+^ signals were analyzed using the AQuA data type presets (Supplementary Material Table 2). ROIs were drawn around the entire cell territory and around the cell soma to obtain Ca^2+^ events from single astrocytes. “Draw anterior” was selected to insert an arrow facing toward the top of the field of view, to indicate true north for analysis. For input in the Detection Pipeline box, we used the parameters shown in Supplementary Material Table 3. In the Export window, Events and features and Movies with overlay were selected before selecting Export/Save. For AquA analysis, we defined Non-Propagating events as those with a traveling distance of 0.99 μm or below, while calcium events with a propagation distance of greater than 1 μm were considered as Propagating events.

### Sampling and statistics

Statistical analyses were performed in Origin Lab, using the Shaprio-Wilk test to test data for normality. Normally distributed datasets were compared using two-sample t-tests. Non-normally distributed datasets were tested using Mann-Whittney for unpaired data. A p-value less than 0.5 was used to determine statistically significant. Unless otherwise indicated, data were expressed as mean ± SEM. Figure legends contain sample sizes and statistical tests used for each experiment.

## Acknowledgements

We thank Grace Hall for assistance with Figure 1B. The authors acknowledge the assistance of the Integrated Microscopy and Imaging Laboratory at the Texas A&M College of Medicine. RRID:SCR_021637. This work was funded by a research grant from the National Institutes of Health (NIH)/NINDS, R21 NS121959 to MR and RS. AM and DA were partially funded by NIH T32 GM135115 and NIH T32 GM135748 grants respectively.

## Contents of Supplementary Material

1 word document and 4 movies

## Author contributions

DAA and AM performed all experiments, analyzed all the data and created the figures. BLS performed experiments and analyzed data. MP, ET, CM, GB, AB and AT analyzed data. MR and RS conceived the project, trained and supervised DAA, AM, BLS, MP, ET, CM, GB, AB and AT, coordinated and designed experiments, and provided resources and funding. MR and RS wrote the manuscript along with DAA and AM, and all authors contributed to editing the final version of the manuscript.

## Data Availability Statement

The data that support the findings of this study are available from the corresponding authors upon reasonable request.

## Supplementary Material For

**Supplementary Material Figure 1.**
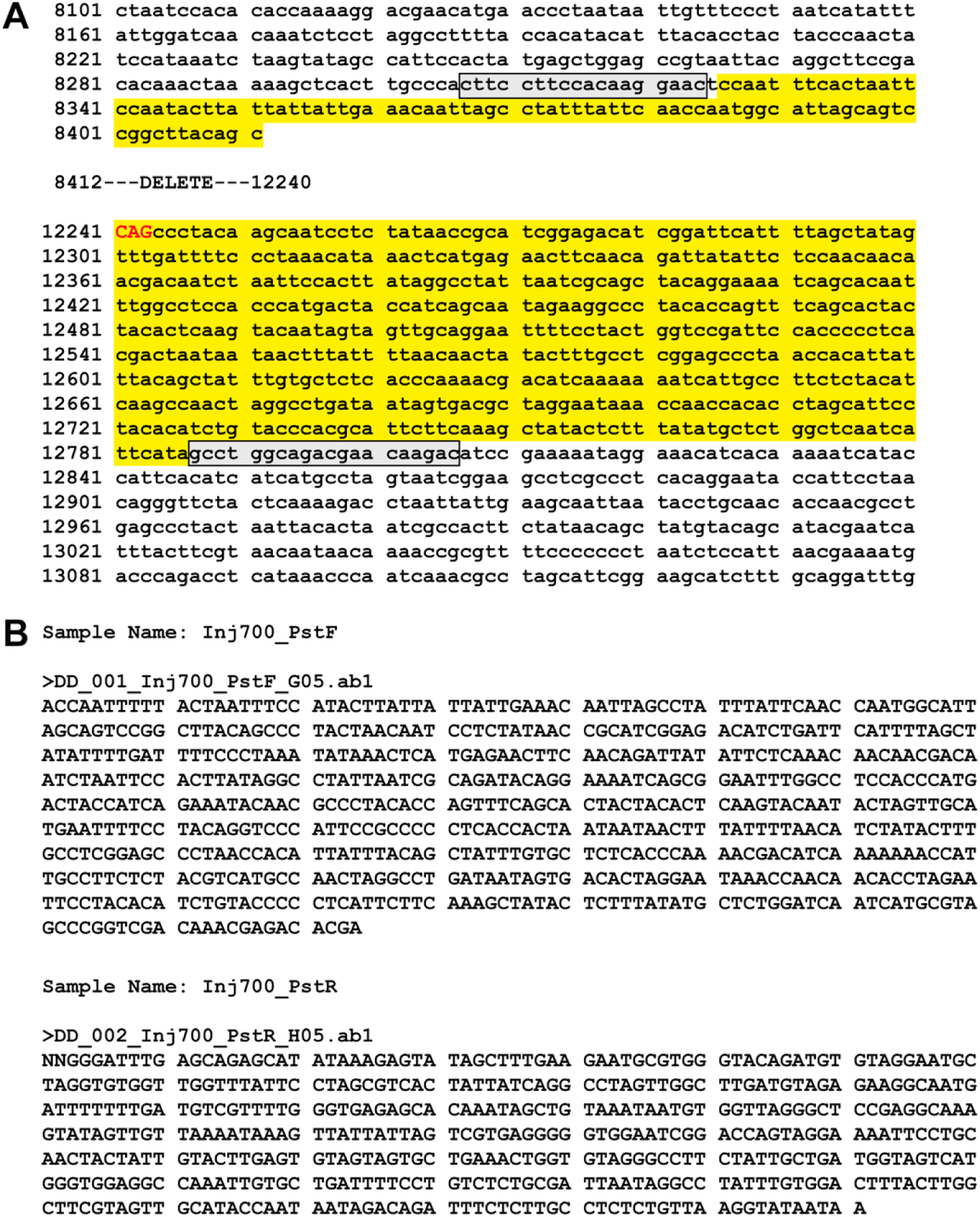
(A) Sequence alignment between mouse mtDNA genome (NC_005089.1) and ∼700bp nested PCR-amplified fragment from PstI-treated DLS (Fig 1E). Numbers to the left indicate nucleotide positions in mitochondrial genome. Gray boxed sequence: Forward and Reverse nested primers. Regions of sequence overlap are highlighted in yellow. Capitalized red bases are the remnants of a PstI site. A 3828 bp deleted region is indicated between overlapping (yellow) regions. **(B)** Forward (top) and reverse (bottom) raw sequences for the ∼700bp nested PCR-amplified fragment from PstI-treated DLS (Fig 1E), used to generate the alignment in (A).

**Supplementary Material Table 1.**
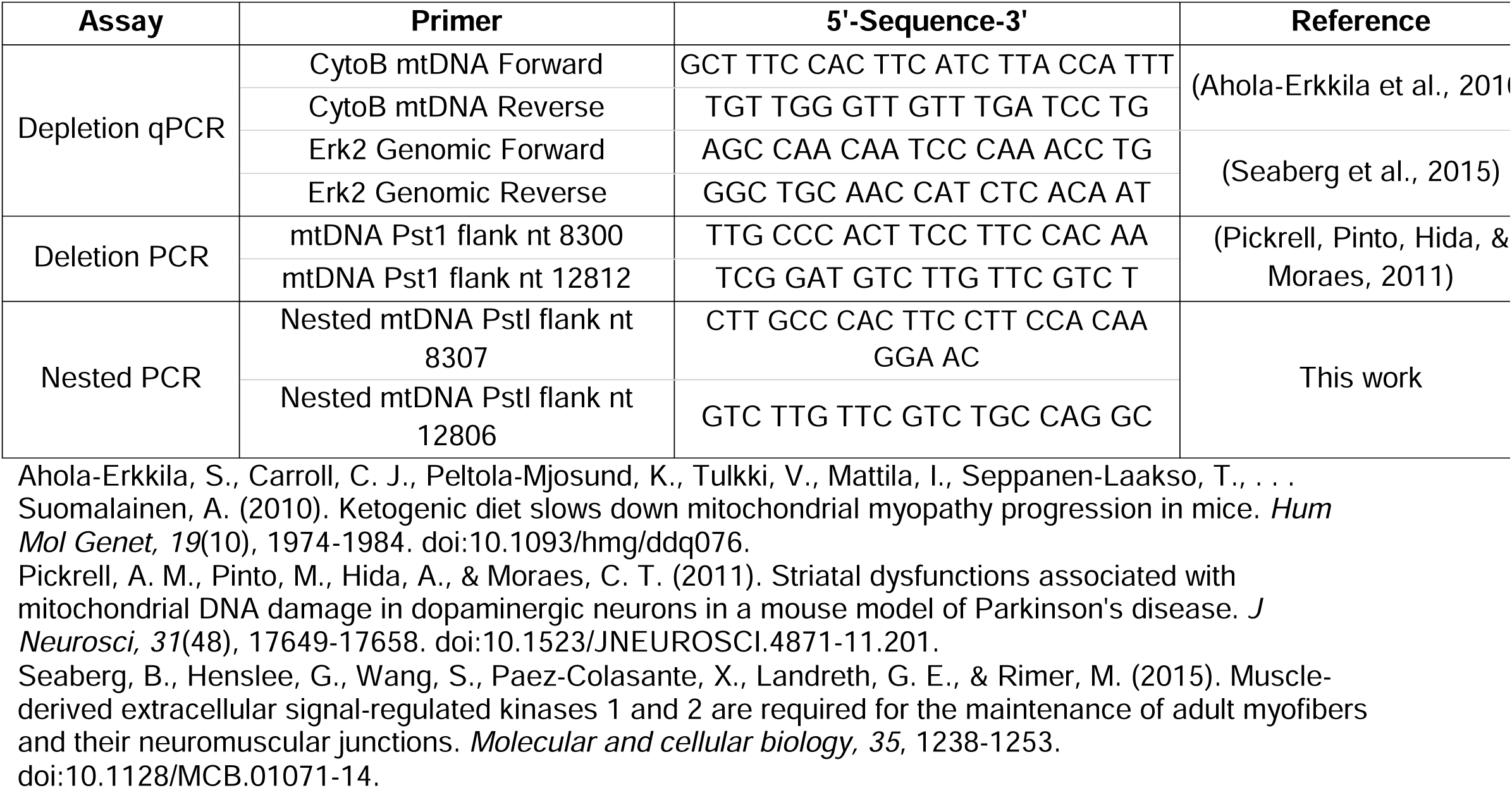
DNA primers used in this study.

**Supplementary Material Table 2.** AQuA presets parameters.

| **AQuA data type** | **Pre-set parameters** |
| --- | --- |
| GCaMP in Vivo | Cyto & Lck |
| Temporal resolution | 0.5 seconds/frame |
| Spatial resolution | 0.5 $\mu\text{m}$/pixel |
| Exclude pixels shorter than | 5 pixels to border |

**Supplementary Material Table 3.**
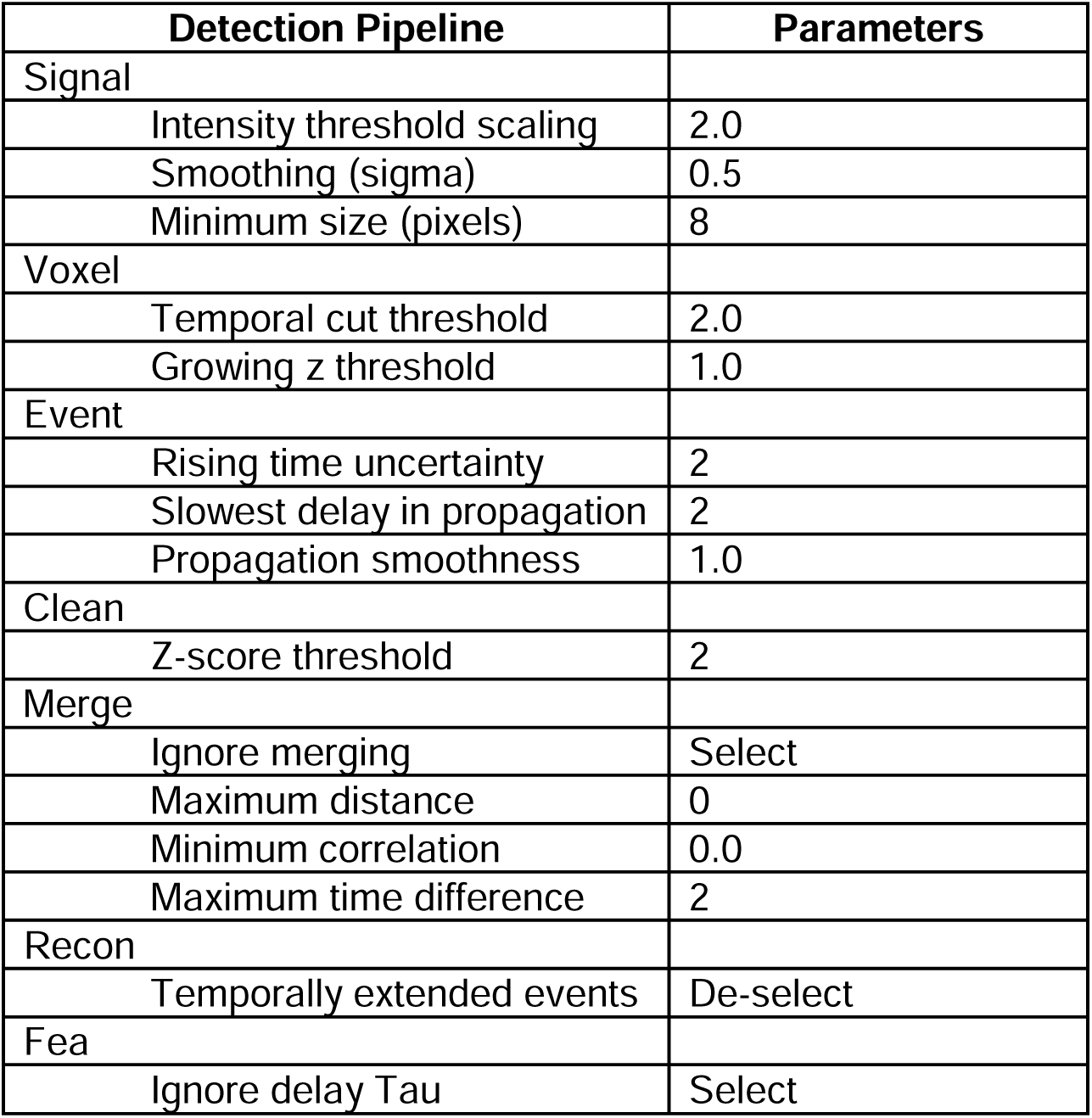
AQuA detection pipeline box parameters.

| **Detection Pipeline** | **Parameters** |
| --- | --- |
| Signal |  |
| Intensity threshold scaling | 2.0 |
| Smoothing (sigma) | 0.5 |
| Minimum size (pixels) | 8 |
| Voxel |  |
| Temporal cut threshold | 2.0 |
| Growing z threshold | 1.0 |
| Event |  |
| Rising time uncertainty | 2 |
| Slowest delay in propagation | 2 |
| Propagation smoothness | 1.0 |
| Clean |  |
| Z-score threshold | 2 |
| Merge |  |
| Ignore merging | Select |
| Maximum distance | 0 |
| Minimum correlation | 0.0 |
| Maximum time difference | 2 |
| Recon |  |
| Temporally extended events | De-select |
| Fea |  |
| Ignore delay Tau | Select |

## Supplementary Material Movie Legends

**Supplemental Movie 1.** Mito-GCaMP expressing astrocytic mitochondria in the HPC with spontaneous Ca^2+^ influx events. Non-Propagating events are pseudo-colored in green and Propagating events are pseudo-colored in red.

**Supplemental Movie 2.** Mito-GCaMP+PstI expressing astrocytic mitochondria in the HPC with spontaneous Ca^2+^ influx events. Non-Propagating events are pseudo-colored in green and Propagating events are pseudo-colored in red.

**Supplemental Movie 3.** Mito-GCaMP expressing astrocytic mitochondria in the DLS with spontaneous Ca^2+^ influx events. Non-Propagating events are pseudo-colored in green and Propagating events are pseudo-colored in red.

**Supplemental Movie 4.** Mito-GCaMP+PstI expressing astrocytic mitochondria in the DLS with spontaneous Ca^2+^ influx events. Non-Propagating events are pseudo-colored in green and Propagating events are pseudo-colored in red.

## REFERENCES

1. Bacman, S. R., Williams, S. L., Pinto, M., & Moraes, C. T. (2014). The use of mitochondria-targeted endonucleases to manipulate mtDNA. Methods Enzymol, 547, 373–397. doi:10.1016/B978-0-12-801415-8.00018-7

2. Batiuk, M. Y., Martirosyan, A., Wahis, J., de Vin, F., Marneffe, C., Kusserow, C., … Holt, M. G. (2020). Identification of region-specific astrocyte subtypes at single cell resolution. Nat Commun, 11(1), 1220. doi:10.1038/s41467-019-14198-8

3. Bazargani, N., & Attwell, D. (2016). Astrocyte calcium signaling: the third wave. Nature Neuroscience, 19(2), 182–189. doi:10.1038/nn.4201

4. Escartin, C., Galea, E., Lakatos, A., O’Callaghan, J. P., Petzold, G. C., Serrano-Pozo, A., … Verkhratsky, A. (2021). Reactive astrocyte nomenclature, definitions, and future directions. Nat Neurosci, 24(3), 312–325. doi:10.1038/s41593-020-00783-4

5. Fecher, C., Trovo, L., Muller, S. A., Snaidero, N., Wettmarshausen, J., Heink, S., … Misgeld, T. (2019). Cell-type-specific profiling of brain mitochondria reveals functional and molecular diversity. Nat Neurosci, 22(10), 1731–1742. doi:10.1038/s41593-019-0479-z

6. Fukui, H., & Moraes, C. T. (2009). Mechanisms of formation and accumulation of mitochondrial DNA deletions in aging neurons. Hum Mol Genet, 18(6), 1028–1036. doi:10.1093/hmg/ddn437

7. G□bel, J., Engelhardt, E., Pelzer, P., Sakthivelu, V., Jahn, H. M., Jevtic, M., … Bergami, M. (2020). Mitochondria-Endoplasmic Reticulum Contacts in Reactive Astrocytes Promote Vascular Remodeling. Cell Metab, 31(4), 791–808 e798. doi:10.1016/j.cmet.2020.03.005

8. Harwig, M. C., Viana, M. P., Egner, J. M., Harwig, J. J., Widlansky, M. E., Rafelski, S. M., & Hill, R. B. (2018). Methods for imaging mammalian mitochondrial morphology: A prospective on MitoGraph. Anal Biochem, 552, 81–99. doi:10.1016/j.ab.2018.02.022

9. Helassa, N., Podor, B., Fine, A., & Török, K. (2016). Design and mechanistic insight into ultrafast calcium indicators for monitoring intracellular calcium dynamics. Scientific Reports, 6(1), 38276. doi:10.1038/srep38276

10. Huntington, T. E., & Srinivasan, R. (2021). Astrocytic mitochondria in adult mouse brain slices show spontaneous calcium influx events with unique properties. Cell Calcium, 96. doi:10.1101/2020.06.21.163618

11. Ignatenko, O., Chilov, D., Paetau, I., de Miguel, E., Jackson, C. B., Capin, G., … Suomalainen, A. (2018). Loss of mtDNA activates astrocytes and leads to spongiotic encephalopathy. Nat Commun, 9(1), 70. doi:10.1038/s41467-017-01859-9

12. Jackson, J. G., & Robinson, M. B. (2018). Regulation of mitochondrial dynamics in astrocytes: Mechanisms, consequences, and unknowns. Glia, 66(6), 1213–1234. doi:10.1002/glia.23252

13. John Lin, C. C., Yu, K., Hatcher, A., Huang, T. W., Lee, H. K., Carlson, J., … Deneen, B. (2017). Identification of diverse astrocyte populations and their malignant analogs. Nat Neurosci, 20(3), 396–405. doi:10.1038/nn.4493

14. Khakh, B. S., & Deneen, B. (2019). The Emerging Nature of Astrocyte Diversity. Annu Rev Neurosci, 42, 187–207. doi:10.1146/annurev-neuro-070918-050443

15. Lee, S., & Min, K. T. (2018). The Interface Between ER and Mitochondria: Molecular Compositions and Functions. Mol Cells, 41(12), 1000–1007. doi:10.14348/molcells.2018.0438

16. Lee, Y., Messing, A., Su, M., & Brenner, M. (2008). GFAP promoter elements required for region-specific and astrocyte-specific expression. Glia, 56(5), 481–493. doi:10.1002/glia.20622

17. Paolicelli, R. C., Sierra, A., Stevens, B., Tremblay, M. E., Aguzzi, A., Ajami, B., … Wyss-Coray, T. (2022). Microglia states and nomenclature: A field at its crossroads. Neuron, 110(21), 3458–3483. doi:10.1016/j.neuron.2022.10.020

18. Pickrell, A. M., Pinto, M., Hida, A., & Moraes, C. T. (2011). Striatal dysfunctions associated with mitochondrial DNA damage in dopaminergic neurons in a mouse model of Parkinson’s disease. J Neurosci, 31(48), 17649–17658. doi:10.1523/JNEUROSCI.4871-11.2011

19. Rong, Z., Tu, P., Xu, P., Sun, Y., Yu, F., Tu, N., … Yang, Y. (2021). The Mitochondrial Response to DNA Damage. Front Cell Dev Biol, 9, 669379. doi:10.3389/fcell.2021.669379

20. Srivastava, S., & Moraes, C. T. (2005). Double-strand breaks of mouse muscle mtDNA promote large deletions similar to multiple mtDNA deletions in humans. Hum Mol Genet, 14(7), 893–902. doi:10.1093/hmg/ddi082

21. Stephen, T. L., Gupta-Agarwal, S., & Kittler, J. T. (2014). Mitochondrial dynamics in astrocytes. Biochem Soc Trans, 42(5), 1302–1310. doi:10.1042/BST20140195

22. Stogsdill, J. A., Harwell, C. C., & Goldman, S. A. (2023). Astrocytes as master modulators of neural networks: Synaptic functions and disease-associated dysfunction of astrocytes. Ann N Y Acad Sci, 1525(1), 41–60. doi:10.1111/nyas.15004

23. Venegas, V., Wang, J., Dimmock, D., & Wong, L. J. (2011). Real-time quantitative PCR analysis of mitochondrial DNA content. *Curr Protoc Hum Genet*, Chapter 19, Unit 19 17. doi:10.1002/0471142905.hg1907s68

24. Wang, Y., DelRosso, N. V., Vaidyanathan, T. V., Cahill, M. K., Reitman, M. E., Pittolo, S., … Poskanzer, K. E. (2019). Accurate quantification of astrocyte and neurotransmitter fluorescence dynamics for single-cell and population-level physiology. Nat Neurosci, 22(11), 1936–1944. doi:10.1038/s41593-019-0492-2

25. Yang, Z., & Wang, K. K. (2015). Glial fibrillary acidic protein: from intermediate filament assembly and gliosis to neurobiomarker. Trends Neurosci, 38(6), 364–374. doi:10.1016/j.tins.2015.04.003

26. Youle, R. J., & van der Bliek, A. M. (2012). Mitochondrial fission, fusion, and stress. Science, 337(6098), 1062–1065. doi:10.1126/science.1219855

27. Zimmer, T. S., Orr, A. L., & Orr, A. G. (2024). Astrocytes in selective vulnerability to neurodegenerative disease. Trends Neurosci, 47(4), 289–302. doi:10.1016/j.tins.2024.02.008

